# Transformation of alignment files improves performance of variant callers for long-read RNA sequencing data

**DOI:** 10.1101/2022.02.08.479579

**Authors:** Vladimir B. C. de Souza, Ben T. Jordan, Elizabeth Tseng, Elizabeth A. Nelson, Karen K. Hirschi, Gloria Sheynkman, Mark D. Robinson

**Affiliations:** Department of Molecular Life Sciences and SIB Swiss Institute of Bioinformatics, University of Zurich, 8057 Zurich, Switzerland; Department of Molecular Physiology and Biological Physics, University of Virginia School of Medicine, Charlottesville, VA 22903, USA; PacBio, San Francisco, USA; Department of Cell Biology and Cardiovascular Research Center, University of Virginia School of Medicine, Charlottesville, VA 22908, USA; Department of Medicine, Yale University School of Medicine; Department of Genetics, Yale University School of Medicine; Yale Cardiovascular Research Center, Yale University School of Medicine, New Haven, CT 06511, USA; Center for Public Health Genomics and UVA Cancer Center, University of Virginia, Charlottesville, VA 22908, USA

## Abstract

Long-read RNA sequencing (lrRNA-seq) produces detailed information about full-length transcripts, including novel and sample-specific isoforms. Furthermore, there is opportunity to call variants directly from lrRNA-seq data. However, most state-of-the-art variant callers have been developed for genomic DNA. Here, there are two objectives: first, we perform a mini-benchmark on GATK, DeepVariant, Clair3, and NanoCaller primarily on PacBio Iso-Seq, data, but also on Nanopore and Illumina RNA-seq data; second, we propose a pipeline to process spliced-alignment files, making them suitable for variant calling with DNA-based callers. With such manipulations, high calling performance can be achieved using DeepVariant on Iso-seq data.

## Background

The detection of genetic variants from next-generation sequencing (NGS) data remains of high interest for applications in clinical diagnostics and to improve our understanding of genetic diseases [1–3]. The most popular variant detection tools have been developed for short-read DNA sequencing data, including GATK [4], bcftools [5], FreeBayes [6] and Platypus [7], among others. However, since the short reads are typically not long enough to encompass multiple variants in a single read, they cannot be phased, *i*.*e*., co-associated to individual isoforms. Fortunately, with the increase in throughput and accuracy of long-read technologies, opportunities for detection of genetic variants from long reads are expanding. For example, IsoPhase [8] was developed to call and phase SNPs from PacBio long-read RNA-seq (lrRNA-seq) data, *i*.*e*. Iso-Seq data, though it does not characterize insertions or deletions (indels). Such information linked to full-length reads offers the opportunity to predict open reading frames (ORFs) with variations that alter protein coding potential [9,10] or transcriptional outcomes, including frame shifts (from indels), truncations or extensions (from altered stop codons), and disrupted splice sites [11]. However, such information is not incorporated in protein prediction. For example, when SQANTI [9] predicts ORFs from long reads, SNPs and indels are reverted to the sequence of the reference genome, losing potentially important patient-specific variations.

Several tools have been designed for calling variants from long reads of DNA aligned to a reference genome, including: DeepVariant [12], Clair3 [13], NanoCaller [14] (for SNP/indel calling); Longshot [15] (for SNP calling); PEPPER-Margin-DeepVariant [16] (for SNP/indel calling from nanopore sequencing data); pbsv [17] (for structural variant calling); and WhatsHap [18] (for variant phasing). Calling variants from lrRNA-seq alignments is also of high interest. For example, TAMA [19] calls variants directly from long reads aligned to a reference genome. Reference-free isoform clustering strategies exist, including IsoCon [20], where a “polishing” step is done to correct errors while keeping variants. Nevertheless, since isoform-clustering and reference-alignment approaches operate at per-isoform and per-gene coverage, respectively, isoform-level approaches tend to show lower sensitivity.

Here, we focus on calling genetic variants from lrRNA-seq reads. Specifically, we perform a mini-benchmark and incorporate existing tools that call variants from DNA-seq or short-read RNA-seq data. The GATK pipeline has already been repurposed to call SNPs and indels from short-read RNA-seq data by using the function SplitNCigarReads (SNCR) [4], which removes intronic regions from mapped reads. DeepVariant, Clair3 and NanoCaller use a deep learning (DL) approach in which variants are detected by analysis of read-alignment images; Clair3 uses a pileup model to call most variants, and a more computationally-intensive full-alignment model to handle more complex variants. All the DL-based tools have been trained and tested on DNA sequencing reads, but not on lrRNA-seq data. In this work, we compare the performance of GATK, DeepVariant, Clair3 and NanoCaller to call variants from Iso-Seq data and comparisons to Nanopore and Illumina RNA-seq are made. We identify factors that influence variant calling performance, including read coverage, proximity to splice junctions, presence of homopolymers, and allele-specific expression. Finally, we present a pipeline to manipulate spliced alignments of BAM files, such that files are suitable for variant calling with DeepVariant and Clair3.

## Results and Discussion

To call variants from lrRNA-seq alignments, we found that transformations of the BAM alignment encodings are critical. This is because while variant calling from aligned DNA sequences data involves analysis of contiguously aligned reads, variant calling from lrRNA-seq alignments must handle reads with gaps representing intronic regions. For example, GATK employs the SNCR function to split reads at introns (Ns in their CIGAR string), thus converting a single isoform alignment into a set of reads representing distinct exons (Fig. 1A). However, SNCR also applies the primary-alignment flag to only one of the split reads and all others receive a supplementary-alignment flag, which can affect performance of downstream tools. Thus, we developed a tool, flagCorrection, to ensure all fragments retain the original flag (Fig. 1A; Additional file 1: Fig. S1 shows an IGV screenshot).

**Fig. 1.**
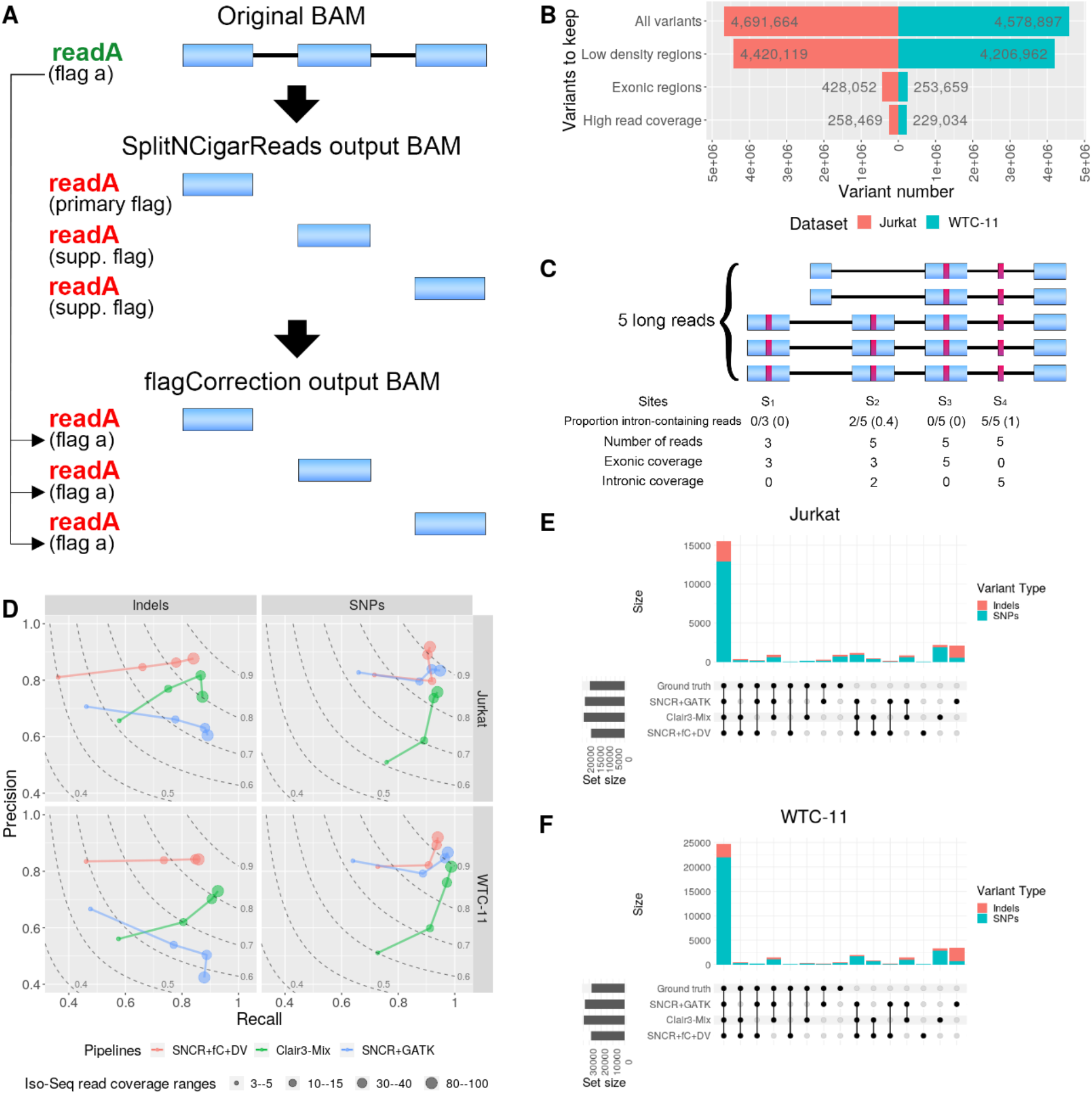
Alignment file transformation for optimised calling of genetic variants from lrRNA-seq data and variant calling performance across the best pipelines on PacBio Iso-Seq reference datasets. (**A**) Alignment file (BAM) transformations to make spliced lrRNA-seq alignments suitable for variant calling. First, GATK’s SNCR function is used to split the reads at Ns in their cigar string, such that exons become distinct reads. Second, the flagCorrection function attributes the flag of the original read to all corresponding fragment reads. (**B**) The number of genetic variants kept in the ground-truth (Illumina DNA-seq) variant call format (VCF) files (for Jurkat and WTC-11 datasets) after filtering; y-axis refers to variant sites that are successively retained, as follows: *All variants*, all sites in the VCF files; *Low density regions*, sites residing in regions such that there is a maximum of 3 variants in a 201 bp window; *Exonic regions*, sites where the Iso-Seq coverage is at least 1; *High read coverage*, sites where the short-read coverage is at least 20 and 72 for Jurkat and WTC-11, respectively; see Methods for more details. (**C**) Schematic with proportion of intron-containing reads (N-cigar reads) at four variant sites (red boxes). (**D**) Precision-recall curves; point sizes indicate the filtering ranges for read coverage; dashed lines represent F1-scores. “Clair3-mix” denotes using Clair3 to call SNPs and SNCR+flagCorrection+Clair3 to call indels. SNCR-SplitNCigarReads; fC-flagCorrection; DV-DeepVariant. Additional file 1: Table S1 gives the number of covered true variants in each interval range. (**E**,**F**) UpSet plots show the intersection of variants called by the pipelines with the ground truth for Jurkat (E) and WTC-11 (F) datasets; sites shown here were filtered according to a minimum Iso-Seq read coverage of 20.

To assess the performance of variant callers, we assembled a set of ground-truth variants for two datasets (Jurkat and WTC-11 cell lines) from Illumina DNA-seq data, retaining only variants from high confidence regions (see Methods) and for which there is sufficient corresponding lrRNA-seq coverage (numbers after filtering are shown in Fig. 1B; we use the term “coverage” to refer to exonic coverage throughout this paper). Our primary analysis here is based on PacBio HiFi reads, while results for Nanopore and Illumina RNA-seq variant calling are presented in the Supplement.

We first evaluated the performance of DeepVariant on Iso-Seq data. To measure the performance gains of DeepVariant calls from manipulated BAM files (Fig. 1A), we called variants from Jurkat and WTC-11 Iso-Seq datasets using three variations: DeepVariant with no alignment manipulation, DeepVariant combined with SNCR (SNCR+DeepVariant), and DeepVariant combined with both SNCR and flagCorrection (SNCR+flagCorrection+DeepVariant). The precision and recall of each pipeline, separated by variant type (SNP or indel) and across various Iso-Seq read coverage ranges, are shown in Additional file 1: Fig. S2. Generally speaking, SNCR+DeepVariant showed low performance, mainly for SNPs, highlighting the need for flagCorrection. Compared with SNCR+flagCorrection+DeepVariant, DeepVariant with no alignment manipulation showed lower performance when read coverage is low to moderate (≤30-40), mainly because of low recall; however, decent performance of DeepVariant is still achieved when coverage is high (80-100).

We found that variant calling performance can be highly influenced by the presence and proximity of intron-containing alignments (*i*.*e*. alignments that contain Ns in their CIGAR string, which represent introns overlapping the variant site location). Consider Fig. 1C, with five reads that overlap a variant (site S_2_), including three that map to the exonic regions with the variant nucleotides, and two intron-containing reads (N-cigar reads) “split” across the variant. Here, the Iso-Seq read coverage would be three. In such a case, the presence of intron-containing reads that cross the variant region can “spoil” variant calling accuracy. We hypothesize that the cause is the presence of N elements (*i*.*e*., introns) in the CIGAR string, which introduce unknown elements into the images that are used by the DL models.

Conversely, at sites with similar exonic coverage (*i*.*e*., Iso-Seq read coverage) but without intron-containing reads (*e*.*g*., site S_1_), variant calling performance tends to be higher. We note also the high proportion of intron-containing reads occurs more frequently in lower-coverage regions (Additional file 1: Fig. S3; Fig. 1C, sites S_2_ and S_4_), as intron-containing reads do not count for read coverage at a site. Taken together, this explains why DeepVariant with no alignment manipulation showed a closer performance to SNCR+flagCorrection+DeepVariant when coverage is high (*e*.*g*., 80-100; see also Additional file 1: Fig. S2), which typically involve cases of exonic regions and hence, SNCR+flagCorrection makes little difference. More directly, Additional file 1: Fig. S4 shows the recall of DeepVariant with no alignment manipulation is heavily dependent on the proportion of intron-containing reads, with extremely low recall when this proportion is high (0.67-1). However, correct management (SNCR+flagCorrection) of split reads and alignment flags mitigates this issue, allowing for DeepVariant to maintain a high performance.

Next, we evaluated the performance of Clair3. Similarly to the DeepVariant comparisons, we compared Clair3’s performance from unmodified BAMs to those subjected to SNCR and/or flagCorrection. Additional file 1: Fig. S5 shows the performance of Clair3-based pipelines using variants merged from both pileup and full-alignment models (recommended by Clair3’s developers [13]), and Additional file 1: Fig. S6 shows variants called only by the pileup model. These results show that the full-alignment model could not accurately call variants from lrRNA-seq data. Moreover, although SNCR+flagCorrection+Clair3 on the pileup model increases indel calling precision while maintaining recall compared to Clair3 with no alignment manipulation (Additional file 1: Fig. S6), the full-alignment model causes many false positives (FPs). Thus, we decided to exclusively use the pileup model to call variants from lrRNA-seq data, and apply this strategy for all subsequent analyses here. Using the pileup model for SNP calling, SNCR+flagCorrection+Clair3 presented a similar precision but slightly lower recall than Clair3. Therefore, we suggest using SNCR+flagCorrection+Clair3 for indels and Clair3 for SNPs, hereafter referred to as “Clair3-mix”.

Using the same strategy as for DeepVariant and Clair3, we compared the performance of NanoCaller-based pipelines. NanoCaller and SNCR+flagCorrection+NanoCaller generally failed to call variants from Iso-Seq data (recall approximately to zero); despite SNCR+NanoCaller showing a higher recall, it is still much lower compared to the other pipelines (Additional file 1: Fig. S7) and is therefore left out of subsequent comparisons.

To compare the performance of DeepVariant and Clair3 with SNCR+GATK, we selected the most accurate version of their pipelines found so far. Precision-recall curves are shown in Fig. 1D, split by variant type (indel and SNP). For indel calling, SNCR+flagCorrection+DeepVariant and Clair3-mix were the best pipelines (similar F1 scores; see Additional file 1: Table S2). However, the DeepVariant-based pipeline showed higher precision, while the recall of Clair3-mix and SNCR+GATK was slightly higher.

SNCR+GATK showed lower precision to call indels. For SNP calling, all three pipelines showed similar performance at high coverage (80-100 Iso-Seq reads), but Clair3-mix showed lower precision at lower coverage (≤10-15 Iso-Seq reads). Taken together, SNCR+flagCorrection+DeepVariant was the best performing single pipeline. Intersections of the called variants compared to the ground truth are shown in Fig. 1E-F. Notably, most of the true variants (Jurkat, 85%; WTC-11, 90%) were called by all pipelines (true positives, TPs); a considerable number of variants were called by Clair3-mix and/or SNCR+GATK, but were absent from the ground truth (FPs); most FPs from SNCR+GATK are indels.

Next, we tested DeepVariant, Clair3 and GATK, with or without BAM manipulation, on Nanopore lrRNA-seq data (from WTC-11 cells; see Methods). Additional file 1: Fig. S8A shows the precision and recall of these pipelines by read-coverage ranges, highlighting that Nanopore lrRNA-seq performance was generally lower than PacBio Iso-Seq, potentially due to the higher error rate of reads. Additional file 1: Fig. S8B shows the intersection of called variants with the ground truth, underscoring the relatively high number of FP indels from Nanopore data. As shown in Additional file 1: Fig. S8C, systematic errors falsely appear as genetic deletions (various similar examples can be shown in a Zenodo repository [21], file name *igv_screenshots_nanopore_alignments*.*zip*).

Although variants called from short reads cannot be phased, which was our original motivation to study variant calling on lrRNA-seq data, we next called variants from Illumina short-read RNA-seq for comparison. We used the same pipelines tested on lrRNA-seq (GATK and DeepVariant-based pipelines), except the Clair3-based pipelines (Clair3’s documentation does not recommend its use with short reads). Moreover, DeepVariant on Illumina RNA-seq without any BAM manipulation was extremely slow (process was killed after 3 weeks using 40 cores; with BAM manipulation, it took around 1.5 hours).

Additional file 1: Fig. S9A shows the performance of the tested pipelines on Illumina RNA-seq, highlighting that SNP calling performance was similar to Iso-Seq (read coverage 30-40), but required a higher read coverage (400-500). In contrast, indel calling was more challenging for Illumina RNA-seq data, becoming worse when coverage is higher than 10-15 for SNCR+flagCorrection+Deepvariant (and higher than 30-40 for SNCR+GATK).

Additional file 1: Fig. S9B shows the intersections of the called variants with the ground truth, and highlights a high occurrence of (indel) FPs, mainly from SNCR+GATK calls. This low performance is partly explained by mapping issues (see various IGV screenshot examples in a Zenodo repository [21], file name *igv_screenshots_illumina_alignments*.*zip*), as short reads are harder to map compared to long reads.

Next, we investigated factors that influence the performance of variant calling specifically from Iso-Seq data. Variants situated close to splice junction boundaries could be more challenging to detect, especially for variant callers that process images of alignments as DeepVariant and Clair3 do. Thus, we determined variant calling performance according to splice junction proximity. Fig. 2A shows the precision, recall, and F1 scores for SNP sites with a minimum Iso-Seq coverage (20 reads or more), separated into those near or not near a splice junction. For a site to be considered near a splice junction, at least half of the reads that contain the site must contain the same splice junction and the site cannot be further than 20 base pairs (bp) away from that junction. For SNP calls near splice junctions, all pipelines showed a drop in recall but a slight increase in precision, indicating that variant calling was more conservative near junctions. However, SNCR+flagCorrection+DeepVariant tended to not detect SNPs near splice junctions, therefore showing a considerable drop in its F1 score. Clair3-mix was the least affected, with no apparent change in its F1 score. On the other hand, to call indels near splice junctions, all pipelines showed a similar drop in their F1 scores (Fig. 2B). Overall, variant calling is less reliable (especially for indel calling) near splice junctions, which could be partially explained by alignment issues near splice junctions due to the presence of these variants (see various IGV screenshot examples in a Zenodo repository [21], file name *igv_screenshots_isoseq_alignments_near_splice_junctions*.*zip*).

**Fig. 2.**
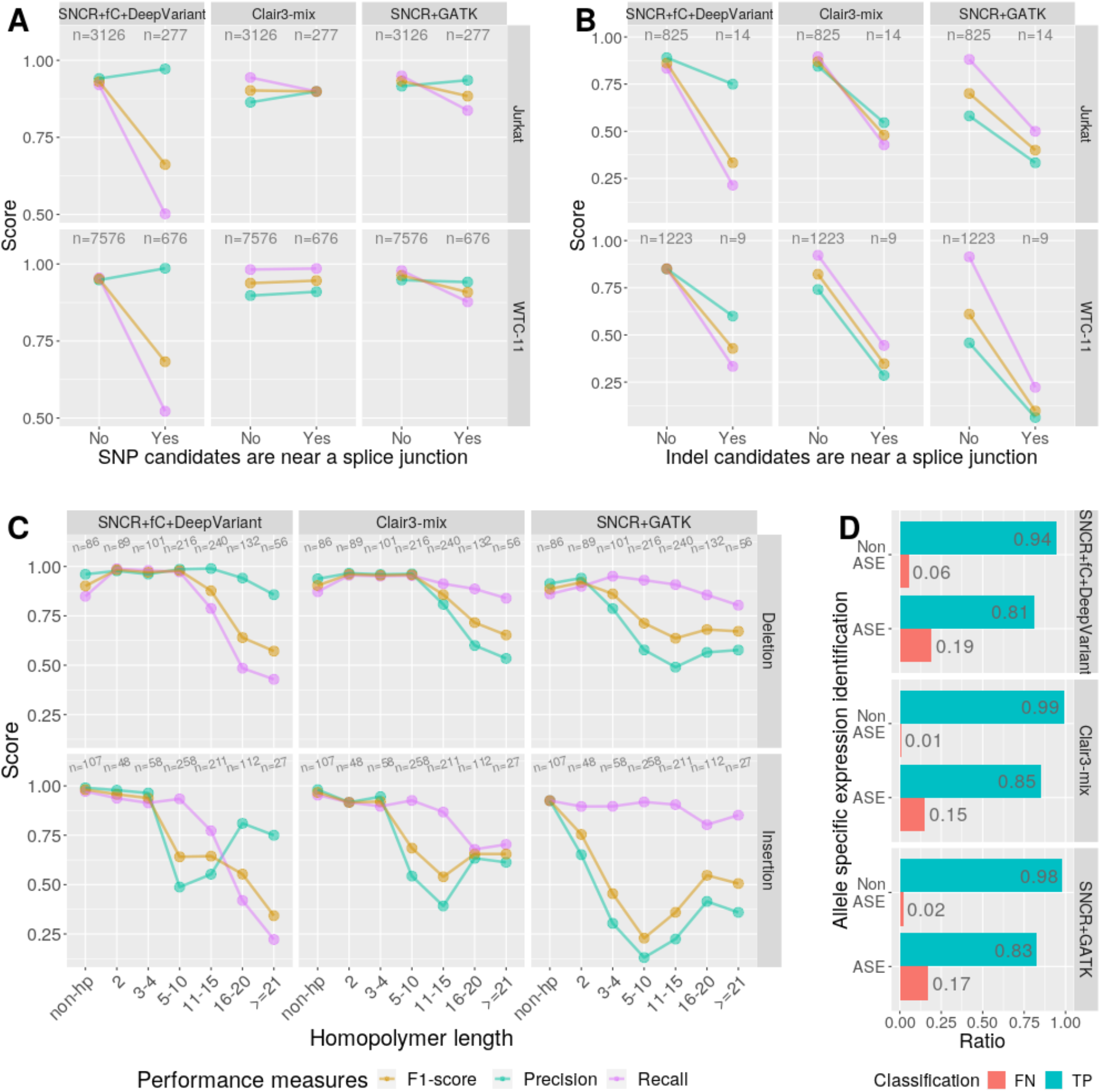
Variant calling performance according to splice junction proximity, homopolymers or allele-specific expression. *n* indicates the number of true variants covered by Iso-Seq data for each calculated performance measure. SNCR-SplitNCigarReads; fC-flagCorrection. In Clair3-based pipelines, only the pileup model was used. Performance measures for SNP (**A**) and indel (**B**) calling of sites far from (*No*) and near to (*Yes*) splice junctions for datasets Jurkat and WTC-11. (**C**) Performance measures of indel calling of sites in non-homopolymers (*non-hp*) and within homopolymers of specified length; results only from WTC-11 dataset. (**D**) FN and TP rates of heterozygous SNP calling from sites in allele-specific expressed (ASE) genes and non-ASE genes; results only from WTC-11 dataset; only sites with a minimum RNA short-read coverage of 40 and minimum Iso-Seq read coverage of 20 were considered.

Another factor that could influence variant calling is the presence of homopolymers. Since sequencing accuracy of long-read platforms is lower in homopolymer-containing regions [22], we evaluated methods to call indels within these regions from the WTC-11 dataset (Jurkat dataset not included due to inadequate read coverage). Fig. 2C shows how precision, recall and F1 score vary according to the length of the homopolymer.

Unsurprisingly, the performance of all pipelines dropped as the length of homopolymer increased; this drop was slightly sharper for insertions.

Since RNA-seq can only observe expressed variants and some genes express only one allele (allele-specific expression; ASE), we hypothesised that variants from ASE genes, corresponding to the lower abundance transcript, would be correlated with a higher false negative (FN; *i*.*e*., an undetected true variant) rate. To investigate this, we used Illumina RNA-seq short reads on WTC-11 cells to categorise heterozygous SNPs in the ground truth set (see Fig. 1B, “High Read Coverage”) as either ASE or non-ASE sites (see Methods).

Fig. 2D highlights that the proportion of FN to TP calls is higher at ASE genes compared to non-ASE genes (chi-squared-test of independence: p-value < 0.001). As expected, for genes expressing a dominant allele, the non-dominant allele would not produce as many reads, and observation of the heterozygous site would be more challenging. Such variants should be considered in future workflows.

## Conclusions

Our mini-benchmark of variant calling from lrRNA-seq data highlights that spliced alignments decrease performance of some standard tools, but after appropriate treatment of alignments and read flags, a high performance can be recovered. In particular, the SplitNCigarReads and flagCorrection functions as applied to input BAM files enable an increase in recall of DeepVariant and the precision of Clair3’s pileup model (for indel calling); Clair3-mix and SNCR+flagCorrection+DeepVariant are among the best-performing pipelines to call indels, the former having higher recall and the latter higher precision. For SNP calling, SNCR+GATK, SNCR+flagCorrection+DeepVariant and Clair3-mix showed similar performance, although Clair3-mix underperformed at lower read coverage. Our results show that when variants are near splice junctions, indel calling was less reliable, and SNCR+flagCorrection+DeepVariant’s recall strongly drops for SNP calling in such regions. Moreover, the performance of all pipelines dropped for indels within homopolymer regions, and we confirmed that ASE genes are a blind spot for RNA-seq-based variant calling.

Overall, we have provided insights on how to call genetic variants from lrRNA-seq data, and we constructed a pipeline (https://github.com/vladimirsouza/lrRNAseqVariantCalling; [23]) for such analyses. This work should be of relevance for applications in genomic medicine, in which variants can be detected and phased directly from lrRNA-seq data collected on patients; phasing tools will need to be demonstrated for the lrRNA-seq setting. It would also be of interest for protein prediction workflows, since genetic variants must be taken into account to correctly predict ORFs and variant protein sequences.

## Methods

### PacBio Iso-Seq datasets

PacBio lrRNA-seq data (*i*.*e*., Iso-Seq) was collected on both Jurkat and WTC-11 cell lines. Jurkat RNA was procured from Ambion (Thermo, PN AM7858) and WTC-11 RNA was extracted from WTC-11 cells (Coriell, GM25256). The RNA was analyzed on a Thermo Nanodrop UV-Vis and an Agilent Bioanalyzer to confirm the RNA concentration and ensure RNA integrity. From the RNA, cDNA was synthesised using the NEB Single Cell/Low Input cDNA Synthesis and Amplification Module (New England Biolabs).

Approximately 300 ng of Jurkat cDNA or WTC-11 cDNA was converted into a SMRTbell library using the Iso-Seq Express Kit SMRT Bell Express Template prep kit 2.0 (Pacific Biosciences). This protocol employs bead-based size selection to remove low mass cDNA, specifically using an 86:100 bead-to-sample ratio (Pronex Beads, Promega). Library preparations were performed in technical duplicate. We sequenced each library on a SMRT cell on the Sequel II system using polymerase v2.1 with a loading concentration of 85 pM. A two-hour extension and 30 hour movie collection time was used for data collection. The ‘ccs‘ command from the PacBio SMRTLink suite (SMRTLink version 9) was used to convert raw reads into Circular Consensus Sequence (CCS) reads. CCS reads with a minimum of three full passes and a 99% minimum predicted accuracy (QV20) were kept for further analysis.

### Nanopore lrRNA-seq dataset

Nanopore lrRNA-seq raw sequences from WTC-11 cells were downloaded from the ENCODE portal [24], identifier ENCFF961HLO.

### Illumina short-read (RNA-seq) datasets

Illumina short reads from WTC-11 cells were downloaded from the NCBI portal [25], identifiers GSM5330767, GSM5330768, and GSM5330769.

### Aligning Iso-Seq lrRNA-seq data to a reference genome

Full-length non-concatemers (FLNC) reads were aligned to the human genome of reference GRCh38.p13 [26] using minimap2 [27] (2.17-r941) with parameters `-ax splice -uf -C5`, and non-primary (secondary, supplementary, and unmapped) alignments were discarded by samtools [28] (1.9); an FLNC-alignment BAM file was generated.

### Aligning Nanopore lrRNA-seq data to a reference genome

Nanopore raw RNA sequences were aligned to the genome of reference GRCh38.p13 using minimap2 with parameters `-ax splice -uf -k14`, and non-primary alignments were discarded by samtools; a Nanopore-lrRNA-seq-alignment BAM file was generated.

### Aligning Illumina RNA-seq data to a reference genome

RNA Illumina short reads were aligned to the genome of reference GRCh38.p13 using STAR (2.7.0f) [31], and secondary and supplementary alignments were discarded by samtools; an Illumina-RNA-seq-alignment BAM file was generated.

### Manipulating BAM files

For each BAM file generated from the alignment sequences of a sequencing technology (PacBio, Nanopore, and Illumina), we used the GATK (4.1.9.0) function SplitNCigarReads (SNCR) to split reads at intronic regions, generating a second BAM file. We generated a third BAM file by correcting flags of the SNCR output BAM with flagCorrection (https://github.com/vladimirsouza/lrRNAseqVariantCalling/blob/main/flagCorrection.r [23]).

### Calling variants from Iso-Seq with DeepVariant

From the flagCorrection output BAM file, genomic variants were called by DeepVariant (1.1.0), using the argument `--model_type PACBIO`. Variants with a QUAL score lower than 15 were filtered out.

### Calling variants from Nanopore lrRNA-seq and Illumina RNA-seq with DeepVariant

Similar to variant calling from Iso-Seq, but using model `--model_type WES`.

### Calling variants from Iso-Seq with Clair3

For SNP calling, from the unmanipulated FLNC-alignment BAM file, variants were called by Clair3 (v0.1-r5), using the argument `--platform=`“`hifi`” and the pre-trained model downloaded from http://www.bio8.cs.hku.hk/clair3/clair3_models/clair3_models.tar.gz, and VCFTools (0.1.16) was used to keep only SNPs. For indel calling, from the flagCorrection output BAM file, Clair3 was run in the same way, and VCFTools was used to keep only indels. In both cases, we considered calls only from the pileup model by using the output file *pileup*.*vcf*.*gz*. The SNP- and indel-only VCF files were concatenated by bcftools (1.9) concat. For sites that culminated with two different variants (one SNP and one indel), we used our function removeRepeatedLowerQualSites.r (https://github.com/vladimirsouza/lrRNAseqVariantCalling/blob/main/tools/removeRepeatedLowerQualSites.r [23]) to remove the variant with the lowest quality (QUAL) value.

### Calling variants from Nanopore lrRNA-seq with Clair3

From appropriate BAM file, variants were called by Clair3, using the argument `--platform=`“`ont`” and the pre-trained model downloaded from http://www.bio8.cs.hku.hk/clair3/clair3_models/r941_prom_hac_g360+g422.tar.gz. We considered calls only from the pileup model by using the output file pileup.vcf.gz.

### Calling variants from (lr)RNA-seq with GATK

From the SNCR output BAM file, read groups were added to the BAM file by Picard [29] AddOrReplaceReadGroups function. Variants were called with the standard GATK pipeline, which consisted of the following steps: generating recalibration table for base quality score recalibration (BQSR) with BaseRecalibrator; applying BQSR with ApplyBQSR; variant calling with HaplotypeCaller; consolidating and genotyping genomic variant call formats (GVCFs) with GenotypeGVCFs; and merging scattered phenotype VCF files with GatherVcfs. For variant-quality score recalibration (VQSR) and filtering, the GATK pipeline used was consisted of the following: VQSR and applying recalibration, both for SNPs and indels, with VariantRecalibrator and ApplyVQSR, respectively.

### Generating the ground truth VCFs for Jurkat and WTC-11 cells

To generate the ground truth of SNPs and indels from Jurkat cells, two Illumina short-read DNA sequencing datasets [30] were downloaded in FASTQ format. The reads from both datasets were aligned to the human reference genome GRCh38.p13 with BWA-MEM [31]. Non-primary (secondary and supplementary) alignments were discarded and the two BAM files were merged by samtools. The same read group was assigned to all reads of the merged BAM by Picard AddOrReplaceReadGroups. Duplicate reads were marked by samtools fixmate followed by samtools markdup. Variants were called with GATK’s pipeline, which consists of: generating recalibration table for base quality score recalibration (BQSR) with BaseRecalibrator; applying BQSR with ApplyBQSR; variant calling with HaplotypeCaller, with ploidy parameter set to diploid; consolidating and genotyping genomic variant call formats (GVCFs) with GenotypeGVCFs; and merging scattered phenotype VCF files with GatherVcfs. For variant-quality score recalibration (VQSR) and filtering, the GATK pipeline used was consisted of the following steps: VQSR and applying recalibration, both for SNPs and indels, with VariantRecalibrator and ApplyVQSR, respectively.

The ground truth variants from WTC-11 cells (a VCF file) was downloaded from the Allen Institute [32]. To generate this VCF, 151 bp paired-end reads, at a mean depth of 100X, were aligned to GRCh38 using BWA-MEM (0.7.13). Duplicates were marked using Picard MarkDuplicates (2.3.0). The GATK’s pipeline (3.5) used consisted of the following steps: local realignment around indels; BQSR; variants calling using HaplotypeCaller; and filtering using VQSR. We kept only variants from chromosomes chr1, …, chr22, chrX, and chrY. Note that both Jurkat and WTC-11 cell lines were derived from male individuals.

### Selecting high confident regions of the ground truth to compare the methods

Since the read coverage of the Jurkat short-read DNA-seq data that we used is not high (overall coverage equal to 38x), only variants residing in regions with short-read coverage higher than 20 reads were considered so as to avoid potential false positives due to low short-read coverage. The variants that passed this coverage filter were considered to be the ground truth for the comparisons of variants called from Iso-Seq Jurkat data.

To avoid mapping/assembly errors (*e*.*g*., due to paralogous or repetitive regions), regions with short-read coverage higher than the 95th-percentile (98 reads) were also ignored. Moreover, to avoid other poorly-aligned regions (*e*.*g*., caused by missing regions of the genome) and after some manual investigation on IGV that highlighted some questionable alignments, any 201 bp window that contains more than three variants was removed. And finally, only regions of the genome that had Iso-Seq coverage >0 were retained.

For the WTC-11 comparisons, a similar strategy was used. But, since the ground-truth VCF file was generated from high-coverage DNA-seq datasets, the arbitrary 20 reads as minimum coverage was not applied. Instead, the 5th-percentile (72 reads) was used as the minimum read coverage. The 95th-percentile (168 reads) was the maximum read coverage.

### Identifying sites that come from ASE genes

Taking the same Illumina-RNA-seq-alignment BAM file (described in “Methods: Aligning Illumina RNA-seq data to a reference genome”) as input, GATK’s ASEReadCounter function was used to calculate read counts per allele of the sites defined by our ground-truth VCF file for WTC-11. We ignored sites with RNA short-read and Iso-Seq coverage lower than 40 and 20 reads, respectively. From the table output by ASEReadCounter, a chi-squared goodness-of-fit test was applied, independently for each site, to test equal frequencies of reference and alternative alleles, and the p-values were corrected by the Benjamini-Hochberg multiple test correction. Sites with q-values lower than 0.05 were considered ASE sites.

### Selecting indels within and outside homopolymer repeats

Sites with Iso-Seq coverage lower than 20 reads were filtered out. To avoid poorly-aligned regions of Iso-Seq reads, any 201 bp window that contains more than three variants (called by any tested pipeline) was removed. To avoid the influence of splice junction proximity, only sites further than 20 bp from any splice junction were considered. To avoid ambiguity in the classification of variant types, heterozygous-alternative variants were filtered out.

## Declarations

### Ethics approval and consent to participate

Ethics approval is not applicable to this work.

### Availability of data and materials

Jurkat and WTC-11 Iso-Seq datasets used in this paper are respectively deposited in the Zenodo repository [33], and NCBI portal, identifier SRR18130587 [34]. WTC-11 Nanopore lrRNA-seq raw reads are available in Encode portal, identifier ENCFF961HLO [35]. WTC-11 Illumina RNA-seq datasets were downloaded from the NCBI portal, identifiers SRR14637256, SRR14637257, and SRR14637258 [36–38]. The human genome, version GRCh38.p13, used here as a reference, is available on the NCBI portal [26]. Illumina DNA-seq datasets from Jurkat cells, used here to compute the Jurkat VCF ground truth, are available in the NCBI portal, identifiers SRR5349449 and SRR5349450 [39,40]. The VCF file used in this paper as the WTC-11 ground truth is available from the Allen Institute [32].

The code used for all analyses presented in this paper, along with flagCorrection function, are available in the public open-source (MIT license) GitHub repository https://github.com/vladimirsouza/lrRNAseqVariantCalling [23] (snapshot in a public Zenodo repository [41]). Important files generated for the analysis are available in a public Zenodo repository [21].

For all benchmarking analyses, several functions were created and are available as an R package in the public open-source (MIT license) GitHub repository https://github.com/vladimirsouza/lrRNAseqBenchmark [42] (snapshot in a public Zenodo repository [43]).

### Authors’ contributions

VBCS, GS, ET, and MDR conceived the study. VBCS designed and implemented all code, ran all data analyses and wrote the manuscript. BJ contributed code and ideas for the analyses. MDR, GS, and ET interpreted data and supervised all analyses. GS, EAN, and KKH conducted experiments and made Jurkat and WTC-11 Iso-Seq data available. All authors revised the manuscript writing and approved the final manuscript.

## Acknowledgements

The long-read sequencing was performed at the Maryland Genomics at the University of Maryland Institute for Genome Sciences. We acknowledge various Robinson lab members for helpful feedback on figures and analyses.

## Supplementary Information

**Additional file 1: Fig. S1** IGV screenshot of three representative BAM files of Iso-Seq reads aligned to the reference genome, in which supplementary alignments are hidden. **Fig. S2** The precision-recall plot of DeepVariant (DV)-based pipelines on Iso-Seq data (PacBio lrRNA-seq), for each dataset (Jurkat or WTC-11), and separated by variant types (indels or SNPs). **Fig. S3** Relationship between the proportion of N-cigar (*i*.*e*., intron-containing) reads and Iso-Seq read coverage (WTC-11 dataset). **Fig. S4** Precision-recall plots of DeepVariant (DV)-based pipelines for variant calling from Iso-Seq data (PacBio lrRNA-seq), according to the proportion of intron-containing (N-cigar) reads (point sizes). **Fig. S5** The precision-recall plot when using both pileup and full-alignment models of Clair3-based pipelines on Iso-Seq data (PacBio lrRNA-seq), for each dataset (Jurkat or WTC-11), and separated by variant types (indels or SNPs). **Fig. S6** The precision-recall plot when using pileup-only model of Clair3-based pipelines on Iso-Seq data (PacBio lrRNA-seq), for each dataset (Jurkat or WTC-11), and separated by variant types (indels or SNPs). **Fig. S7** The precision-recall plot of the SNCR+NanoCaller pipeline on Iso-Seq data (PacBio lrRNA-seq), for each dataset (Jurkat or WTC-11), and separated by variant types (indels or SNPs). **Fig. S8** Variant calling performance on Nanopore lrRNA-seq data. **Fig. S9** Variant calling performance on Illumina RNA-seq data. **Table S1** Number of true indels and SNPs covered by Iso-Seq data, in each read coverage range used in the mini-benchmark, for Jurkat and WTC-11 datasets. **Table S2** Performance measures (precision, recall, and F1 score) of the best tested pipelines (SNCR+flagCorrection+DeepVariant, Clair3-mix, and SNCR+GATK), for each dataset (Jurkat and WTC-11), separated by variant types (indels and SNPs), using different thresholds for minimum Iso-Seq read coverage (Min_coverage). **Table S3** Number of true indels and SNPs covered by Nanopore lrRNA-seq data, in each read coverage range used in the mini-benchmark, for WTC-11 dataset. **Table S4** Number of true indels and SNPs covered by Illumina RNANanopore lrRNA-seq data, in each read coverage range used in the mini-benchmark, for WTC-11 dataset.

