## Supplementary Info for "Transformation of alignment files improves performance of variant callers for long-read RNA sequencing data"

### Additional file 1

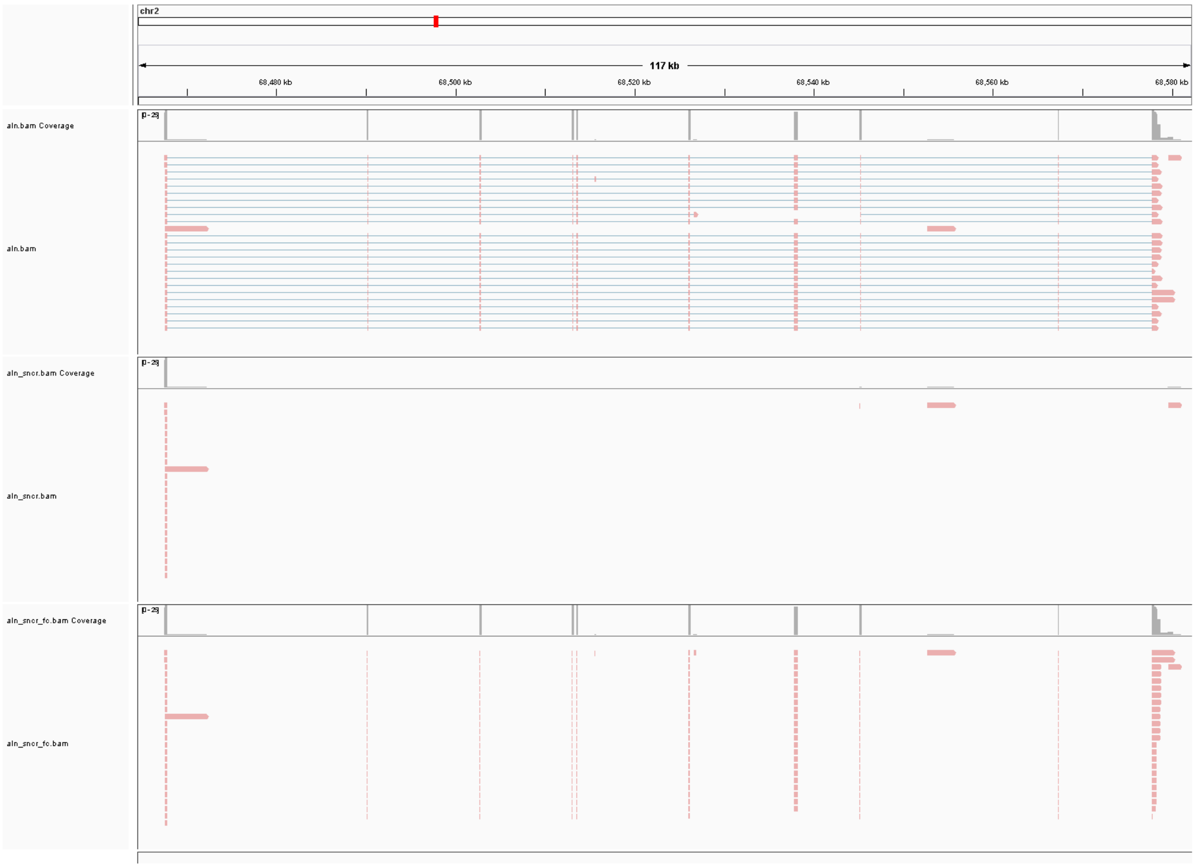


**Fig. S1** IGV screenshot of three representative BAM files of Iso-Seq reads aligned to the reference genome, in which supplementary alignments are hidden. From the top to the bottom, the BAM files are: *aln.bam*, the original read alignments; *aln_sncr.bam*, the SplitNCigarReads (SNCR) output generated from *aln.bam*; *aln_sncr_fc.bam*, the flagCorrection output generated from *aln_sncr.bam*. The tracks named as a BAM file name followed by “Coverage” show the read coverage of the respective BAM file. The pink bars represent exons, while blue lines are introns.

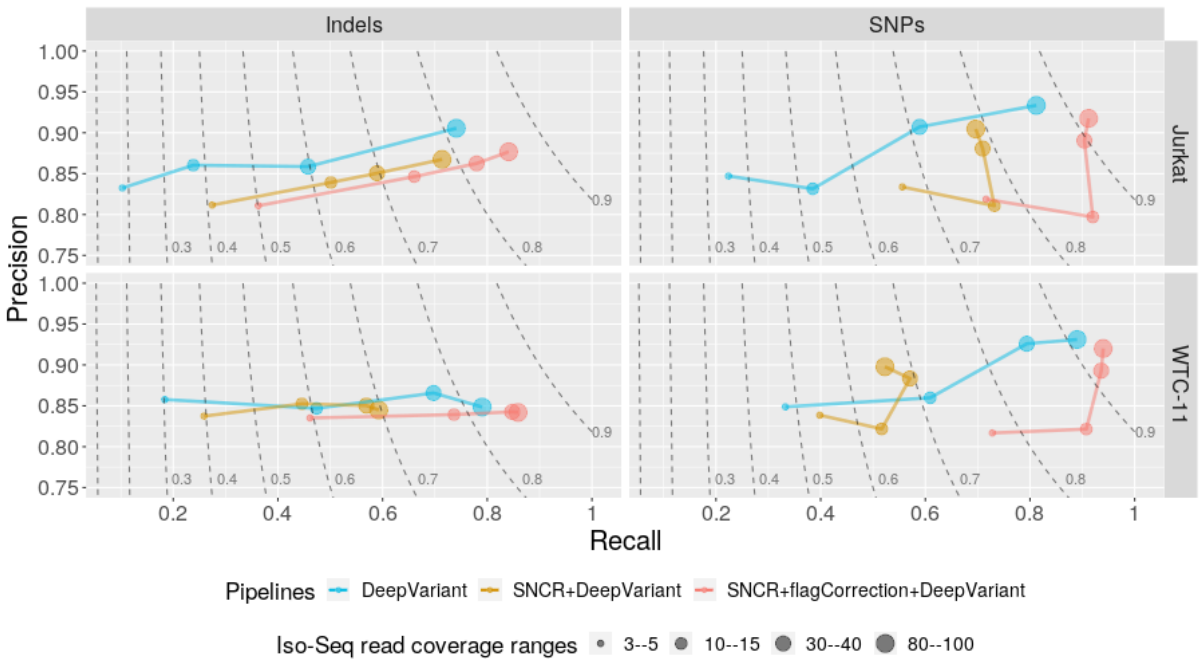


**Fig. S2** The precision-recall plot of DeepVariant (DV)-based pipelines on Iso-Seq data (PacBio lrRNA-seq), for each dataset (Jurkat or WTC-11), and separated by variant types (indels or SNPs). SNCR refers to SplitNCigarReads. Point sizes indicate the filtering ranges for read coverage. Dashed lines are F1-score curves followed by their values. Additional file 1: Table S1 gives the number of covered true variants in each interval range.

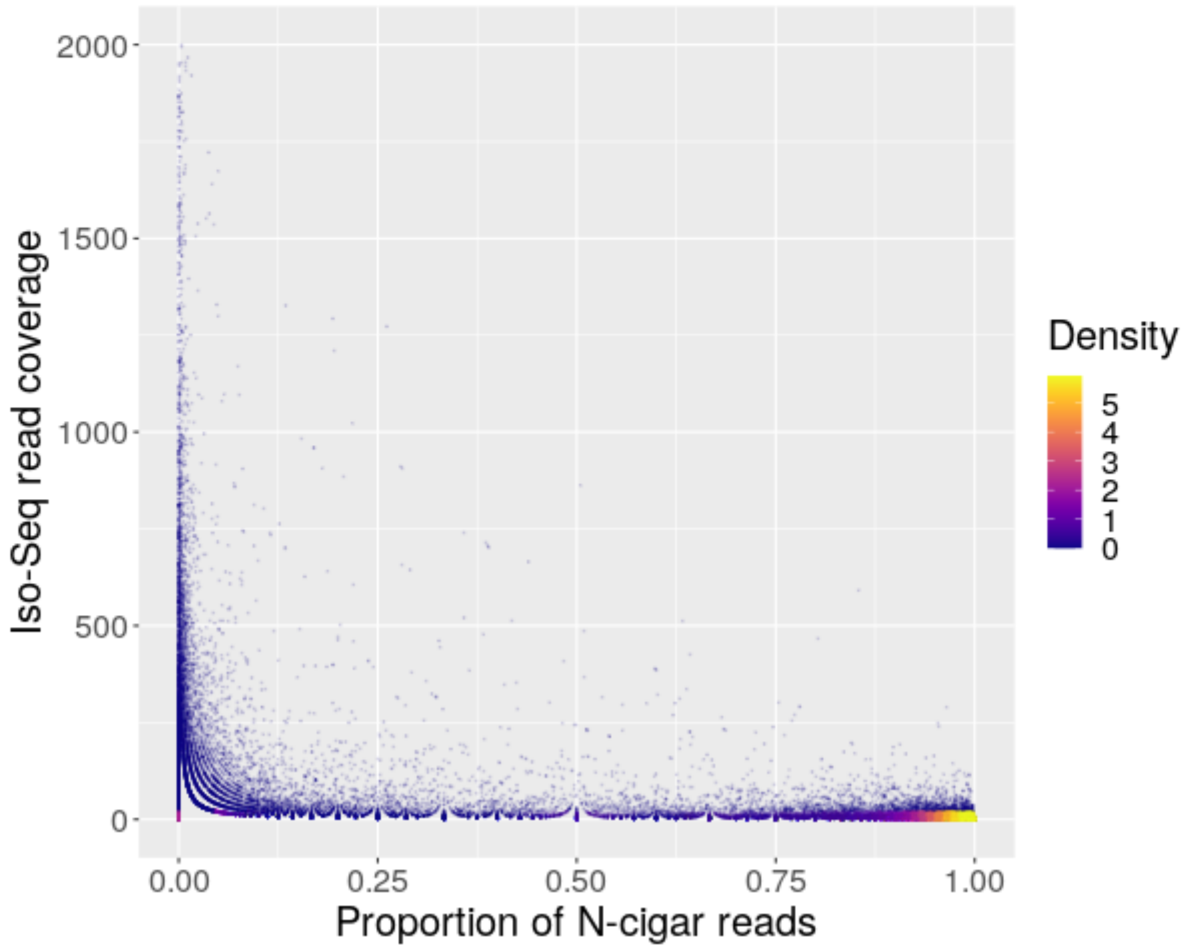


**Fig. S3** Relationship between the proportion of N-cigar (*i.e.*, intron-containing) reads and Iso-Seq read coverage (WTC-11 dataset). N-cigar reads represent reads that contain Ns in their CIGAR string (*i.e.* introns) at the site location (see Fig. 1C for schematic). Each point represents a nucleotide site on the reference genome that is covered by Iso-Seq reads. Colours indicate the density of points. N-cigar reads, which represent introns, are not counted towards the total Iso-Seq read coverage quantification of any given site (y-axis).

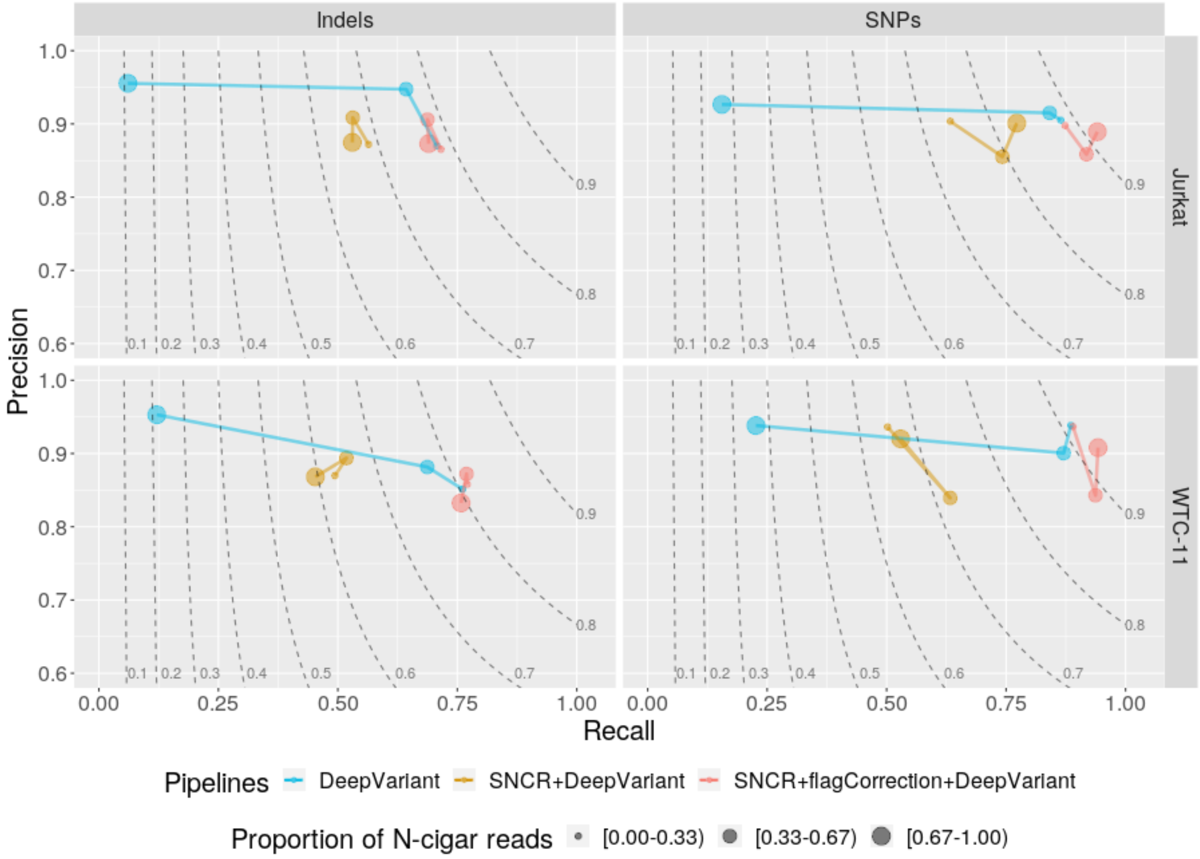


**Fig. S4** Precision-recall plots of DeepVariant (DV)-based pipelines for variant calling from Iso-Seq data (PacBio lrRNA-seq), according to the proportion of intron-containing (N-cigar) reads (point sizes), *i.e.* reads that contain Ns in their CIGAR string at the site location, separated by variant types (indels and SNPs), and dataset (Jurkat and WTC-11). SNCR refers to SplitNCigarReads. In order for the analysis to be less influenced by read coverage, only variant candidate sites with Iso-Seq coverage between 10-20 reads were considered. fC refers to flagCorrection. Number of sites per interval of intron-containing-read proportions: 8,076 [0.00-0.33), 1,636 [0.33-0.7), 6,948 [0.67-1.00). Dashed lines are F1-score curves followed by their values.

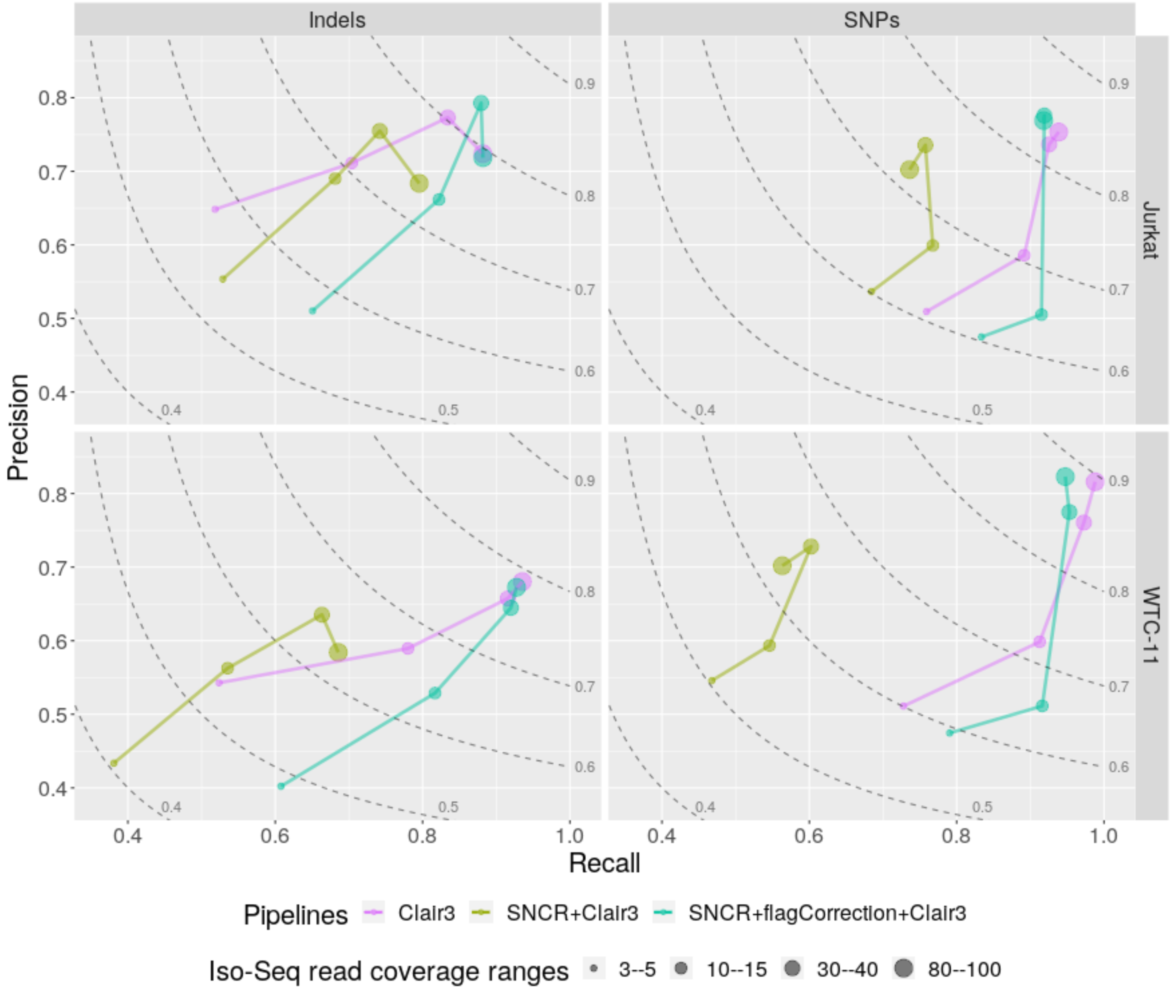


**Fig. S5** The precision-recall plot when using both pileup and full-alignment models of Clair3-based pipelines on Iso-Seq data (PacBio lrRNA-seq), for each dataset (Jurkat or WTC-11), and separated by variant types (indels or SNPs). SNCR refers to SplitNCigarReads. Point sizes indicate the filtering threshold for minimum read coverage. Dashed lines are F1-score curves followed by their values. Additional file 1: Table S1 gives the number of covered true variants in each interval range.

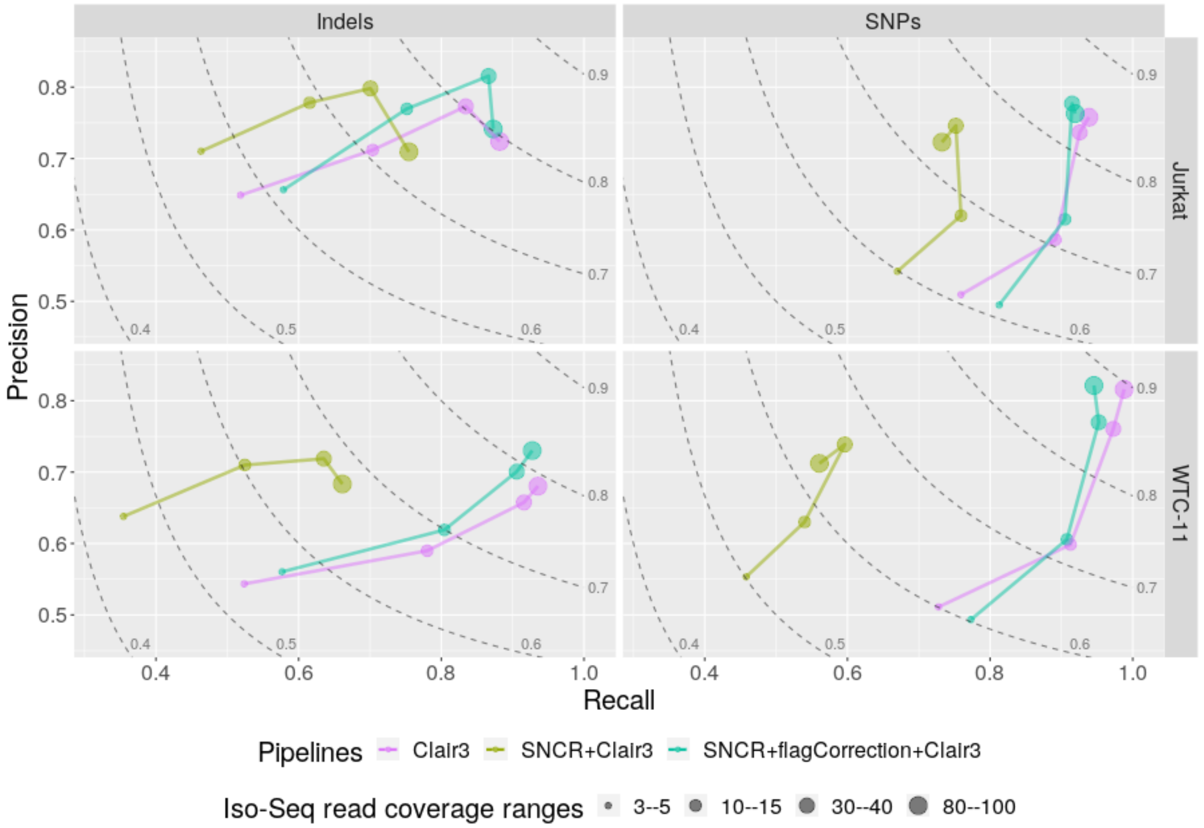


**Fig. S6** The precision-recall plot when using pileup-only model of Clair3-based pipelines on Iso-Seq data (PacBio lrRNA-seq), for each dataset (Jurkat or WTC-11), and separated by variant types (indels or SNPs). SNCR refers to SplitNCigarReads. Point sizes indicate the filtering threshold for minimum read coverage. Dashed lines are F1-score curves followed by their values. Additional file 1: Table S1 gives the number of covered true variants in each interval range.

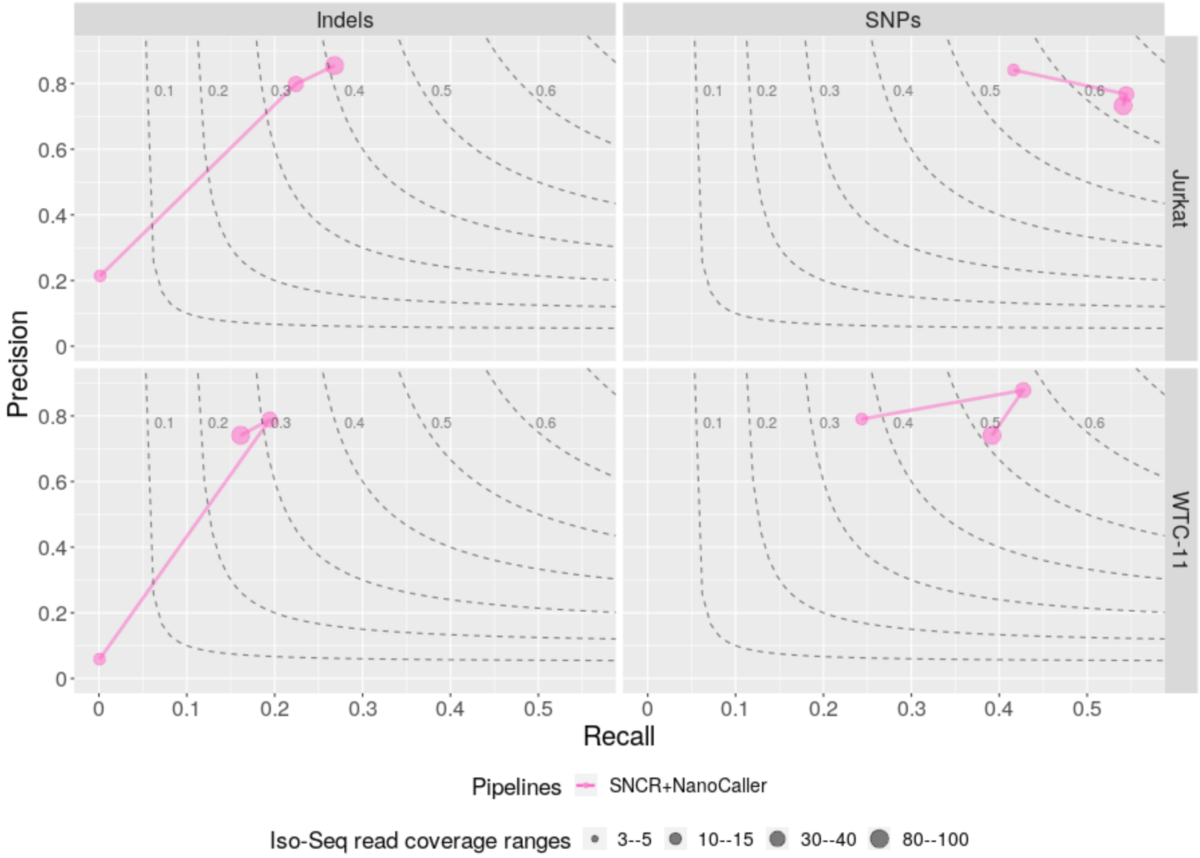


**Fig. S7** The precision-recall plot of the SNCR+NanoCaller pipeline on Iso-Seq data (PacBio lrRNA-seq), for each dataset (Jurkat or WTC-11), and separated by variant types (indels or SNPs). SNCR refers to SplitNCigarReads. Point sizes indicate the filtering threshold for minimum read coverage. Dashed lines are F1-score curves followed by their values. NanoCaller underperforms with SNCR in part because it tries to phase reads based on the called SNPs before calling indels, which is not possible after splitting the exons into reads. Precision and recall are not available when the range of Iso-Seq coverage is 3-5, because SNCR+NanoCaller could not call any variant. Additional file 1: Table S1 gives the number of covered true variants in each interval range.

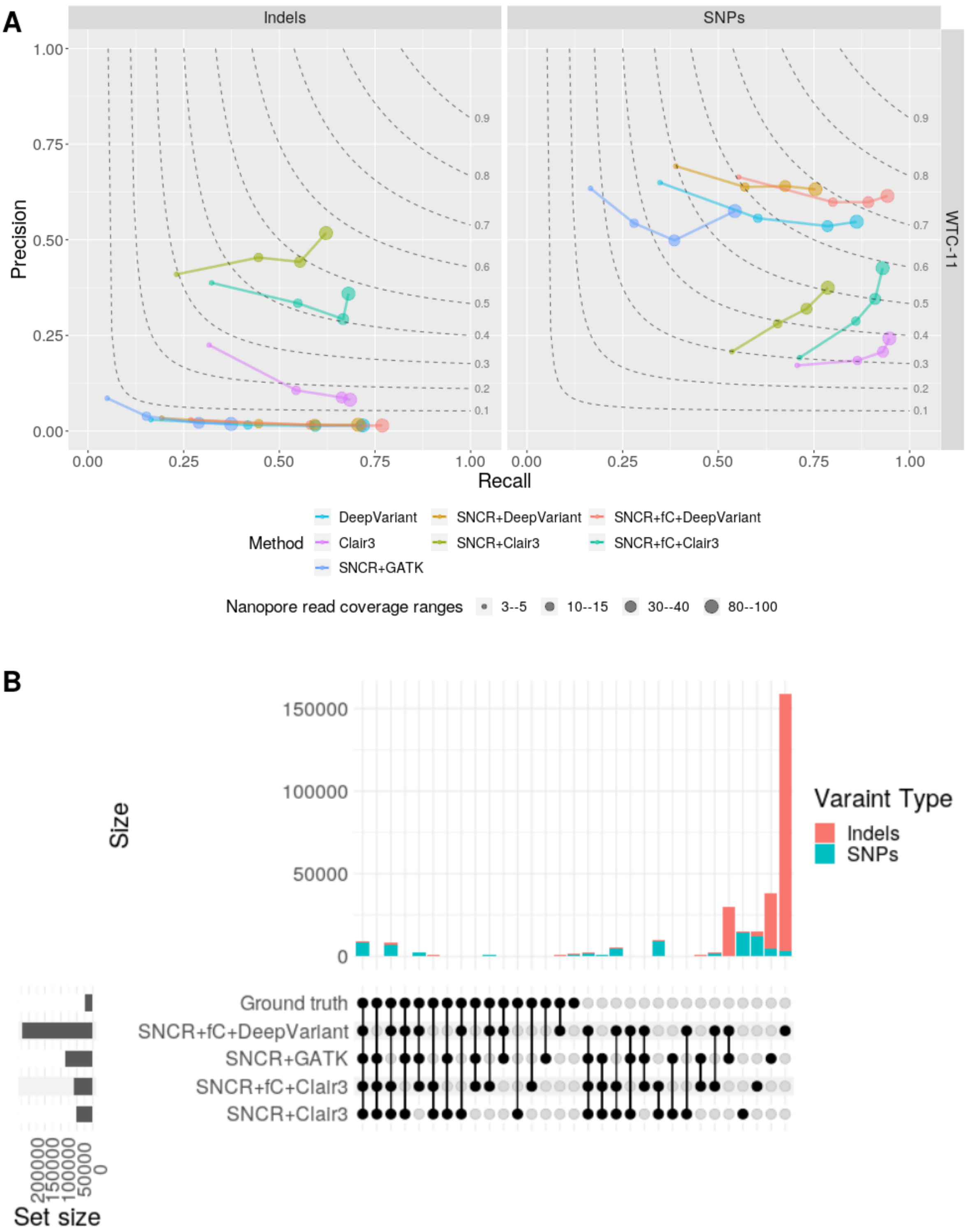
A
Indels
SNPs
WTC-11
Precision
Recall
Method
DeepVariant
Clair3
SNCR+GATK
SNCR+DeepVariant
SNCR+Clair3
SNCR+fC+DeepVariant
SNCR+fC+Clair3
Nanopore read coverage ranges
3--5
10--15
30--40
80--100
B
Size
Varaint Type
Indels
SNPs
Ground truth
SNCR+fC+DeepVariant
SNCR+GATK
SNCR+fC+Clair3
SNCR+Clair3
Set size

C

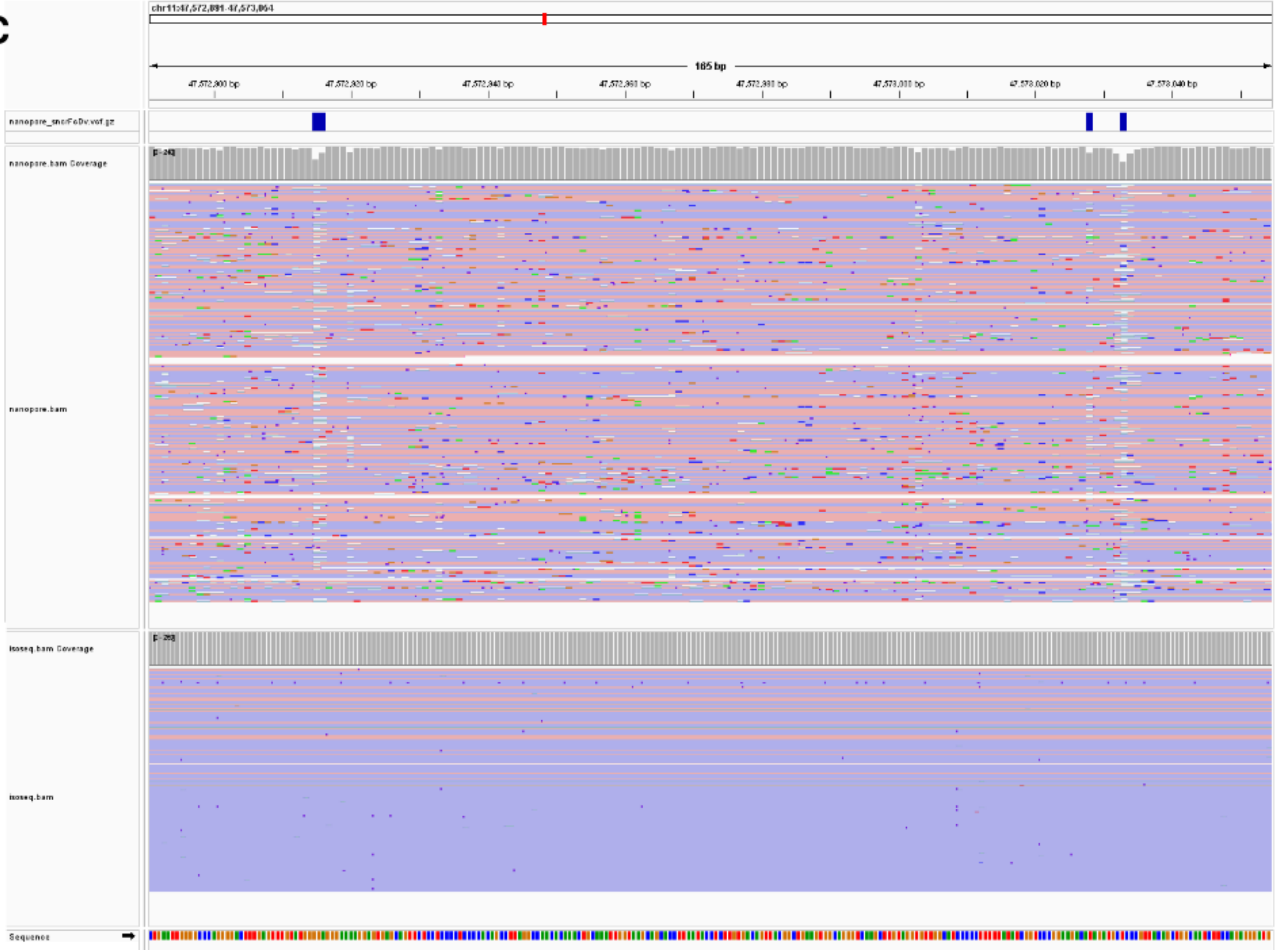

chr11:47,572,891-47,573,064
165 bp
47,572,900 bp
47,572,920 bp
47,572,940 bp
47,572,960 bp
47,572,980 bp
47,573,000 bp
47,573,020 bp
47,573,040 bp
nanopore_sncFoDv.vcf.gz
nanopore.bam Coverage
[0-24]
nanopore.bam
isoseq.bam Coverage
[0-20]
isoseq.bam
Sequence

**Fig. S8** Variant calling performance on Nanopore lrRNA-seq data. (**A**) The precision-recall plot of DeepVariant-based, Clair3-based and GATK pipelines on Nanopore lrRNA-seq data, for WTC-11 dataset, separated by variant type (indels or SNPs). SNCR refers to SplitNCigarReads. Point sizes indicate the filtering ranges for read coverage. Dashed lines are F1-score curves followed by their values. Additional file 1: Table S3 gives the number of covered true variants in each interval range. (**B**) UpSet plot shows the intersection of variants called by the pipelines with the ground truth for WTC-11 datasets; sites shown here were filtered according to a minimum Nanopore read coverage of 20. (**C**) IGV screenshot of false-positive deletion calls at an exonic region. The top track displays the VCF file results, produced by SNCR+flagCorrection+DeepVariant on Nanopore data. Below the VCF track are two representative BAM files, from Nanopore lrRNA-seq (middle) and Iso-Seq (bottom) reads aligned to the reference genome. Grey bars at the top of each BAM file represent respective read coverage. The pink (forward strand) and blue (reverse strand) bars represent the stretch of reads aligned to the exon.

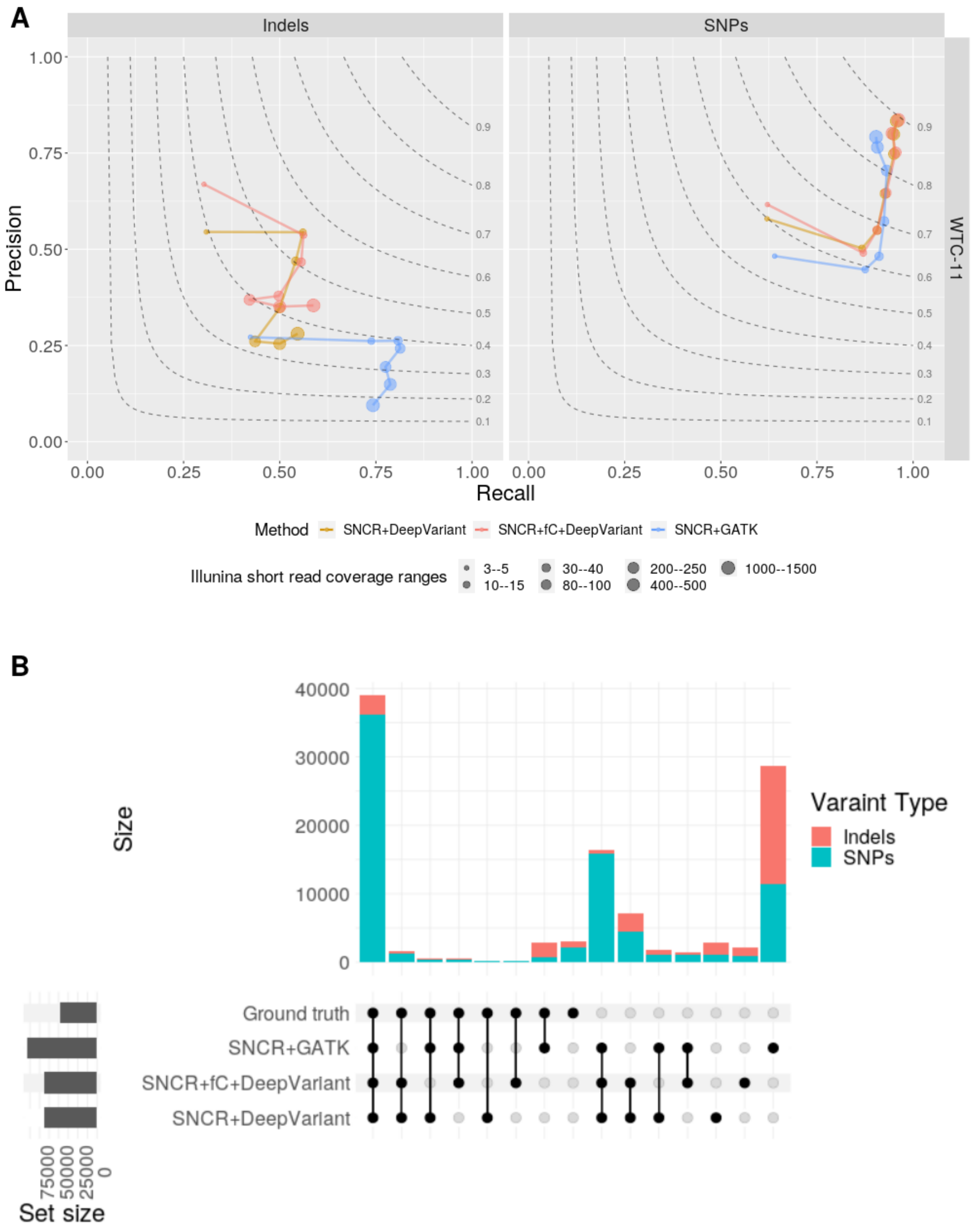


**Fig. S9** Variant calling performance on Illumina RNA-seq data. (**A**) The precision-recall plot of DeepVariant-based (DeepVariant with no alignment manipulation not included) and GATK pipelines on Illumina RNA-seq data, for WTC-11 dataset, and separated by variant types (indels or SNPs). SNCR refers to SplitNCigarReads. Point sizes indicate the filtering ranges for read coverage. Dashed lines are F1-score curves followed by their values. Additional file 1: Table S4 gives the number of covered true variants in each interval range. (**B**) UpSet plot shows the intersection of variants called by the pipelines with the ground truth for WTC-11 datasets; sites shown here were filtered according to a minimum Illumina read coverage of 20.

**Table S1** Number of true indels and SNPs covered by Iso-Seq data, in each read coverage range used in the mini-benchmark, for Jurkat and WTC-11 datasets.

| Dataset | Type | 3-5 | 10-15 | 30-40 | 80-100 |
|---|---|---|---|---|---|
| Jurkat | Indels | 10969 | 2093 | 531 | 220 |
| Jurkat | SNPs | 34352 | 7036 | 1976 | 830 |
| WTC-11 | Indels | 5050 | 1283 | 499 | 248 |
| WTC-11 | SNPs | 24998 | 6957 | 3040 | 1580 |

**Table S2** Performance measures (precision, recall, and F1 score) of the best tested pipelines (SNCR+flagCorrection+DeepVariant, Clair3-mix, and SNCR+GATK), for each dataset (Jurkat and WTC-11), separated by variant types (indels and SNPs), using different thresholds for minimum Iso-Seq read coverage (Min_coverage). SNCR refers to SplitNCigarReads, fC to flagCorrection, and DV to DeepVariant.

| Dataset | Variant_type | Mim_coverage | Pipeline | Precision | Recall | F1-score |
|---|---|---|---|---|---|---|
| Jurkat | Indels | 3 | SNCR+fC+DV | 0.839 | 0.509 | 0.634 |
| Jurkat | Indels | 3 | Clair3-mix | 0.707 | 0.673 | 0.69 |
| Jurkat | Indels | 3 | SNCR+GATK | 0.655 | 0.612 | 0.633 |
| Jurkat | Indels | 15 | SNCR+fC+DV | 0.87 | 0.772 | 0.819 |
| Jurkat | Indels | 15 | Clair3-mix | 0.789 | 0.852 | 0.819 |
| Jurkat | Indels | 15 | SNCR+GATK | 0.6 | 0.854 | 0.704 |
| Jurkat | Indels | 40 | SNCR+fC+DV | 0.871 | 0.8 | 0.834 |
| Jurkat | Indels | 40 | Clair3-mix | 0.79 | 0.871 | 0.828 |
| Jurkat | Indels | 40 | SNCR+GATK | 0.57 | 0.863 | 0.687 |
| Jurkat | Indels | 100 | SNCR+fC+DV | 0.868 | 0.801 | 0.833 |
| Jurkat | Indels | 100 | Clair3-mix | 0.804 | 0.874 | 0.837 |
| Jurkat | Indels | 100 | SNCR+GATK | 0.544 | 0.845 | 0.662 |
| Jurkat | SNPs | 3 | SNCR+fC+DV | 0.832 | 0.809 | 0.821 |
| Jurkat | SNPs | 3 | Clair3-mix | 0.571 | 0.83 | 0.677 |
| Jurkat | SNPs | 3 | SNCR+GATK | 0.827 | 0.775 | 0.8 |
| Jurkat | SNPs | 15 | SNCR+fC+DV | 0.894 | 0.907 | 0.901 |
| Jurkat | SNPs | 15 | Clair3-mix | 0.757 | 0.932 | 0.835 |
| Jurkat | SNPs | 15 | SNCR+GATK | 0.86 | 0.929 | 0.893 |
| Jurkat | SNPs | 40 | SNCR+fC+DV | 0.926 | 0.898 | 0.912 |
| Jurkat | SNPs | 40 | Clair3-mix | 0.839 | 0.939 | 0.886 |
| Jurkat | SNPs | 40 | SNCR+GATK | 0.89 | 0.94 | 0.914 |
| Jurkat | SNPs | 100 | SNCR+fC+DV | 0.947 | 0.892 | 0.919 |
| Jurkat | SNPs | 100 | Clair3-mix | 0.891 | 0.941 | 0.915 |
| Jurkat | SNPs | 100 | SNCR+GATK | 0.915 | 0.941 | 0.928 |
| WTC-11 | Indels | 3 | SNCR+fC+DV | 0.839 | 0.643 | 0.728 |
| WTC-11 | Indels | 3 | Clair3-mix | 0.628 | 0.729 | 0.675 |
| WTC-11 | Indels | 3 | SNCR+GATK | 0.533 | 0.673 | 0.595 |
| WTC-11 | Indels | 15 | SNCR+fC+DV | 0.837 | 0.83 | 0.834 |
| WTC-11 | Indels | 15 | Clair3-mix | 0.709 | 0.893 | 0.791 |
| WTC-11 | Indels | 15 | SNCR+GATK | 0.466 | 0.878 | 0.609 |
| WTC-11 | Indels | 40 | SNCR+fC+DV | 0.838 | 0.854 | 0.846 |
| WTC-11 | Indels | 40 | Clair3-mix | 0.721 | 0.915 | 0.807 |
| WTC-11 | Indels | 40 | SNCR+GATK | 0.44 | 0.895 | 0.59 |
| WTC-11 | Indels | 100 | SNCR+fC+DV | 0.83 | 0.853 | 0.841 |
| WTC-11 | Indels | 100 | Clair3-mix | 0.73 | 0.912 | 0.811 |
| WTC-11 | Indels | 100 | SNCR+GATK | 0.418 | 0.887 | 0.569 |
| WTC-11 | SNPs | 3 | SNCR+fC+DV | 0.847 | 0.845 | 0.846 |

| | | | | | | |
|---|---|---|---|---|---|---|
| WTC-11 | SNPs | 3 | Clair3-mix | 0.619 | 0.859 | 0.72 |
| WTC-11 | SNPs | 3 | SNCR+GATK | 0.841 | 0.814 | 0.827 |
| WTC-11 | SNPs | 15 | SNCR+fC+DV | 0.895 | 0.927 | 0.911 |
| WTC-11 | SNPs | 15 | Clair3-mix | 0.776 | 0.969 | 0.862 |
| WTC-11 | SNPs | 15 | SNCR+GATK | 0.865 | 0.957 | 0.908 |
| WTC-11 | SNPs | 40 | SNCR+fC+DV | 0.911 | 0.932 | 0.922 |
| WTC-11 | SNPs | 40 | Clair3-mix | 0.838 | 0.983 | 0.905 |
| WTC-11 | SNPs | 40 | SNCR+GATK | 0.894 | 0.97 | 0.931 |
| WTC-11 | SNPs | 100 | SNCR+fC+DV | 0.911 | 0.933 | 0.922 |
| WTC-11 | SNPs | 100 | Clair3-mix | 0.867 | 0.988 | 0.923 |
| WTC-11 | SNPs | 100 | SNCR+GATK | 0.915 | 0.971 | 0.942 |

**Table S3** Number of true indels and SNPs covered by Nanopore lrRNA-seq data, in each read coverage range used in the mini-benchmark, for WTC-11 dataset.

| Type | 3-5 | 10-15 | 30-40 | 80-100 |
|---|---|---|---|---|
| Indels | 7829 | 1513 | 525 | 238 |
| SNPs | 35583 | 7199 | 2870 | 1417 |

**Table S4** Number of true indels and SNPs covered by Illumina RNA-seq data, in each read coverage range used in the mini-benchmark, for WTC-11 dataset.

| Type | 3-5 | 10-15 | 30-40 | 80-100 | 200-250 | 400-500 | 1000-1500 |
|---|---|---|---|---|---|---|---|
| Indels | 11848 | 2842 | 842 | 412 | 280 | 160 | 97 |
| SNPs | 62498 | 15840 | 5158 | 2538 | 1772 | 1130 | 779 |
